# Minimal effects of *spargel* (PGC-1α) overexpression in a *Drosophila* mitochondrial disease model

**DOI:** 10.1101/529545

**Authors:** Jack George, Howard T. Jacobs

## Abstract

PGC-1α and its homologues have been proposed to act as master regulators of mitochondrial biogenesis in animals. Most relevant studies have been conducted in mammals, where interpretation is complicated by the fact that there are three partially redundant members of the gene family. In *Drosophila*, only a single PGC-1α homologue, *spargel* (*srl*), is present in the genome. Here we analyzed the effects of *srl* overexpression on phenotype and on gene expression in *tko*^*25t*^, a recessive bang-sensitive mutant with a global defect in oxidative phosphorylation, resulting in a deficiency of mitochondrial protein synthesis. In contrast to previous reports, we found only minimal effects of substantial overexpression of *srl* throughout development, on the expression of a representative set of both mtDNA- and nuclear-encoded OXPHOS-related transcripts, both in *tko*^*25t*^ mutant flies and heterozygous controls. Sex and genetic background appeared to influence the expression of the tested genes, but *srl* overexpression or *tko*^*25t*^ itself did not have clear-cut or systematic effects thereon. These studies provide no support to the concept of spargel as a global regulator of mitochondrial biogenesis.

**Summary blurb:** overexpression of *spargel*, the fly PGC-1α homologue proposed as a mitochondrial biogenesis regulator, has minimal effects on the phenotype of *tko*^*25t*^, considered a fly model for mitochondrial disease

## INTRODUCTION

The PGC1 coactivators are widely considered to be global regulators of bioenergy metabolism, specifically acting to promote mitochondrial biogenesis in many different contexts (Spiegelman, 2007). However, the fact that there are three such factors encoded in mammalian genomes (PGC-1α, PGC-1β and PPRC1, also denoted as PRC) complicates their analysis, due to the combination of tissue or physiological specialization and genetic redundancy (Finck & Kelly, 2006).

In the *Drosophila* genome, a single member of the PGC1 coactivator family, *spargel* (*srl*), is present A *srl* hypomorph, with a P-element promoter insertion, was found to have decreased weight, decreased accumulation of storage nutrients in males, and female sterility (Tiefenböck et al., 2010). In the mutant larval fat body there was decreased respiratory capacity and decreased expression of genes required for mitochondrial biogenesis and activity, with evidence of co-operation with the *Drosophila* NRF-2α homologue Delg, and with responsiveness to insulin signaling. These findings are consistent with spargel acting as a general regulator of mitochondrial biogenesis in the fly. Many subsequent studies have been construed similarly (Mukherjee et al., 2014).

As part of a previous study of phenotypes connected with the *Drosophila* mutant *tko*^*25t*^, we found evidence consistent with a role for spargel in regard to mitochondrial functions (Chen et al., 2012). *tko*^*25t*^ carries a point mutation in the gene encoding mitoribosomal protein S12 (Royden et al., 1987; Shah et al., 1997), which confers larval developmental delay, bang sensitivity, defective male courtship and impaired sound responsiveness (Toivonen et al., 2001). The mutant has an under-representation of mitoribosomal small subunit rRNA and is deficient in all four enzymes of the oxidative phosphorylation (OXPHOS) system that depend on mitochondrial DNA (mtDNA)-encoded subunits (Toivonen et al., 2001; Toivonen et al., 2003). The *tko*^*25t*^ phenotype can be rescued by an additional genomic copy of the mutant *tko* locus (Kemppainen et al., 2009) and partially compensated by altered mtDNA background (Chen et al., 2012) or low sugar diet (Kemppainen et al., 2016).

In our earlier study, flies overexpressing *srl* showed a modest but statistically significant alleviation of the mutant phenotype (Chen et al., 2012). When we later catalogued our strain collection, we concluded that this experiment may have used a strain carrying a genomic duplication of *srl* (designated *srl*^*GR*^, Tiefenböck et al., 2010), rather than the GAL4-dependent *srl* cDNA construct. In order to clarify the effects on *tko*^*25t*^ phenotype of *srl* overexpression at different levels, we proceeded to combine the mutant with different *srl* constructs, having first profiled their effects on expression. In an initial experiment using *srl*^*GR*^, we were able to substantiate the earlier finding, of a modest alleviation of developmental delay. However, this was not upheld in subsequent repeats of the experiment, nor by other strain combinations that overexpress *srl* at a higher level; nor did *srl* overexpression systematically modulate the expression of genes for OXPHOS subunits or the mitochondrial nucleoid protein TFAM. We thus find no consistent evidence to support a role for *srl* in boosting mitochondrial biogenesis in *tko*^*25t*^ flies.

## RESULTS

### *srl* expression in wild-type and *tko*^*25t*^ mutant flies

To assess the effects of the *srl* overexpression in *tko*^*25t*^ mutant flies and heterozygous controls, we first measured the extent of overexpression using qRT-PCR, after combining the relevant chromosomes carrying *srl*^*GR*^, UAS-*srl*, the ubiquitously acting *da*GAL4 driver, the *tko*^*25t*^ mutation and appropriate balancer chromosomes (Fig. 1). To reproduce as closely as possible the previously studied conditions, we created *tko*^*25t*^ flies that were hemizygous for both *srl*^*GR*^ and *da*GAL4, even though there should be no UAS construct present (Fig. 1A). We also analyzed the sexes separately since, in initial trials, we observed a consistently higher endogenous *srl* expression in females than males. Hemizygosity for the *srl*^*GR*^ construct conferred an increase in *srl* RNA in both sexes, proportionate to gene dosage (Fig. 1A). In contrast, UAS-*srl* driven by *da*GAL4 resulted in a more substantial increase in *srl* RNA: ∼four-fold in females and >100-fold in males (Fig. 1B). *srl* RNA was at lower abundance in *tko*^*25t*^ females (though not males: Fig. 1C), and was restored to the wild-type level by *srl*^*GR*^ (Fig. 1D). To test whether increased *srl* RNA due to UAS-*srl* expression was reflected at the protein level, we generated an antibody against spargel, which detected a major band of approximate molecular weight ∼105 kDa and a minor band of ∼125 kDa (Fig. 2A), close to the predicted molecular weight of the protein (118 kDa). These bands were detected in both males and females (Fig. 2B: note that the ∼125 kDa band appears more faintly in females, but is always present at long exposure). The same two bands were detected in S2 cells induced to express V5 epitope-tagged spargel (Fig. 2C, 2D). UAS-*srl* driven by *da*GAL4 led to an increase in detected spargel protein, based on Western blot signal compared with the GAPDH loading control (Fig. 2E, 2F). Note, however, that the increase was proportionately far smaller than that seen at the RNA level, and that the disparity in expression at the RNA level between males and females was not evident in the detected protein.

**Figure 1:**
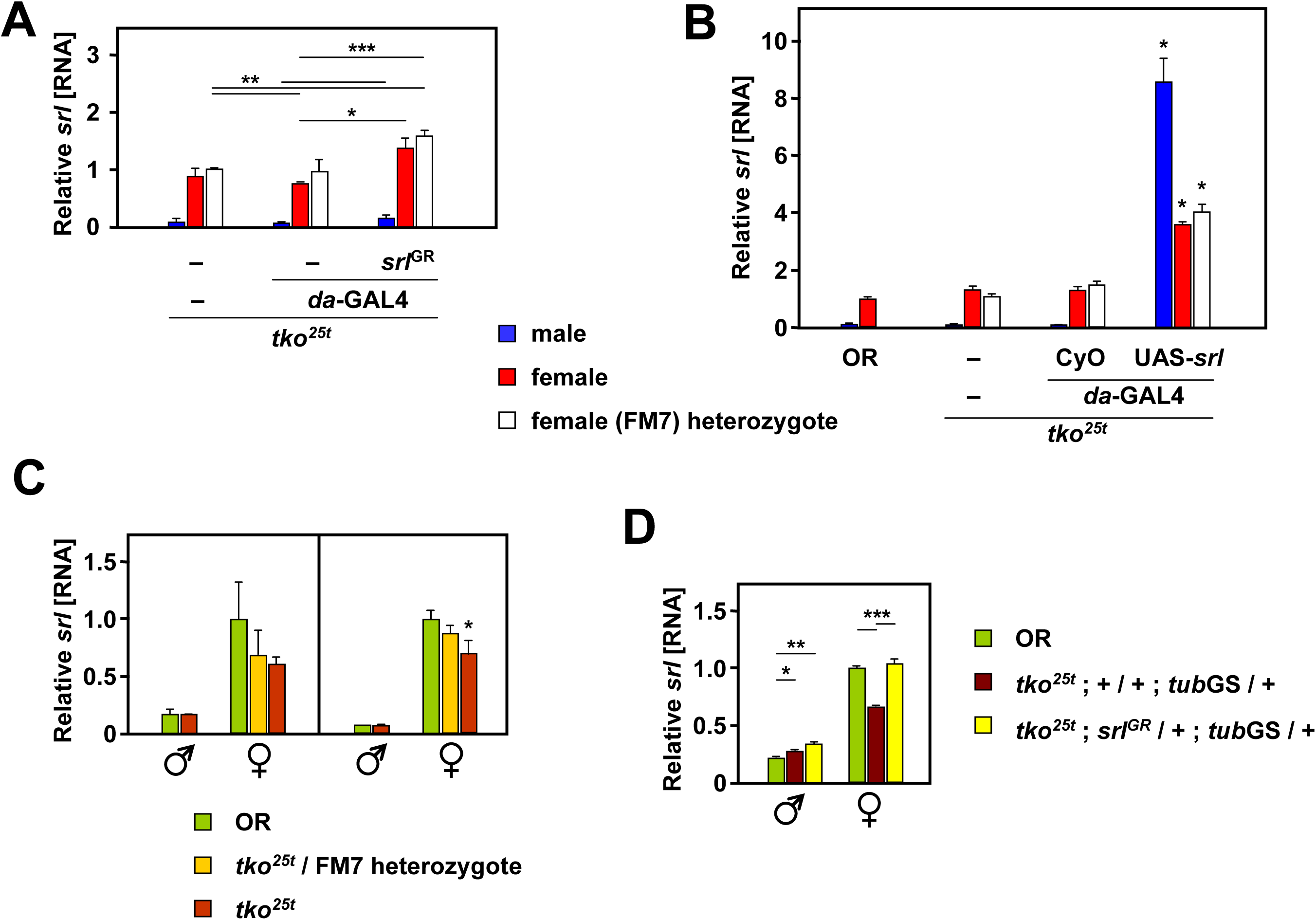
*srl* can be overexpressed by genomic duplication or using the GAL4 system. qRT-PCR measurements of *srl* RNA in adult flies of the indicated genotypes and sex, normalized to values for control females: (A) *tko*^*25t*^ / FM7 heterozygotes, which have a wild-type phenotype; (B, C, D) Oregon R (OR) wild-type. (A) Effect of genomic duplication of *srl* (hemizygosity for *srl*^*GR*^) with or without the additional presence of the *da*GAL4 driver, in the *tko*^*25t*^ background. (B) Effect of UAS-*srl*, with or without the *da*GAL4 driver, in the *tko*^*25t*^ background, alongside OR. Asterisks denote significant differences between flies expressing UAS-*srl* driven by *da*GAL4 and non-expressing controls of the same sex and *tko* genotype (Student’s *t* test, *p* < 0.001). (C) Effect of the *tko*^*25t*^ background (two separate experiments separated by vertical line). Asterisk denotes significant difference from OR flies of the same sex (Student’s *t* test, *p* < 0.05). (D) Combined effect of the *tko*^*25t*^ background and hemizygosity for *srl*^*GR*^. Note that. in this experiment, we substituted the *tub*GS driver background for *da*GAL4, but with similar results. Asterisks denote significant differences, based on Student’s *t* test with Bonferroni correction, comparing flies of a given sex between genotypes: *, **, *** – p < 0.05, 0.01, 0,.001, respectively. Note that all statistical analyses are based on ddCT values from the qRT-PCR data, not the fold differences as plotted.

**Figure 2:**
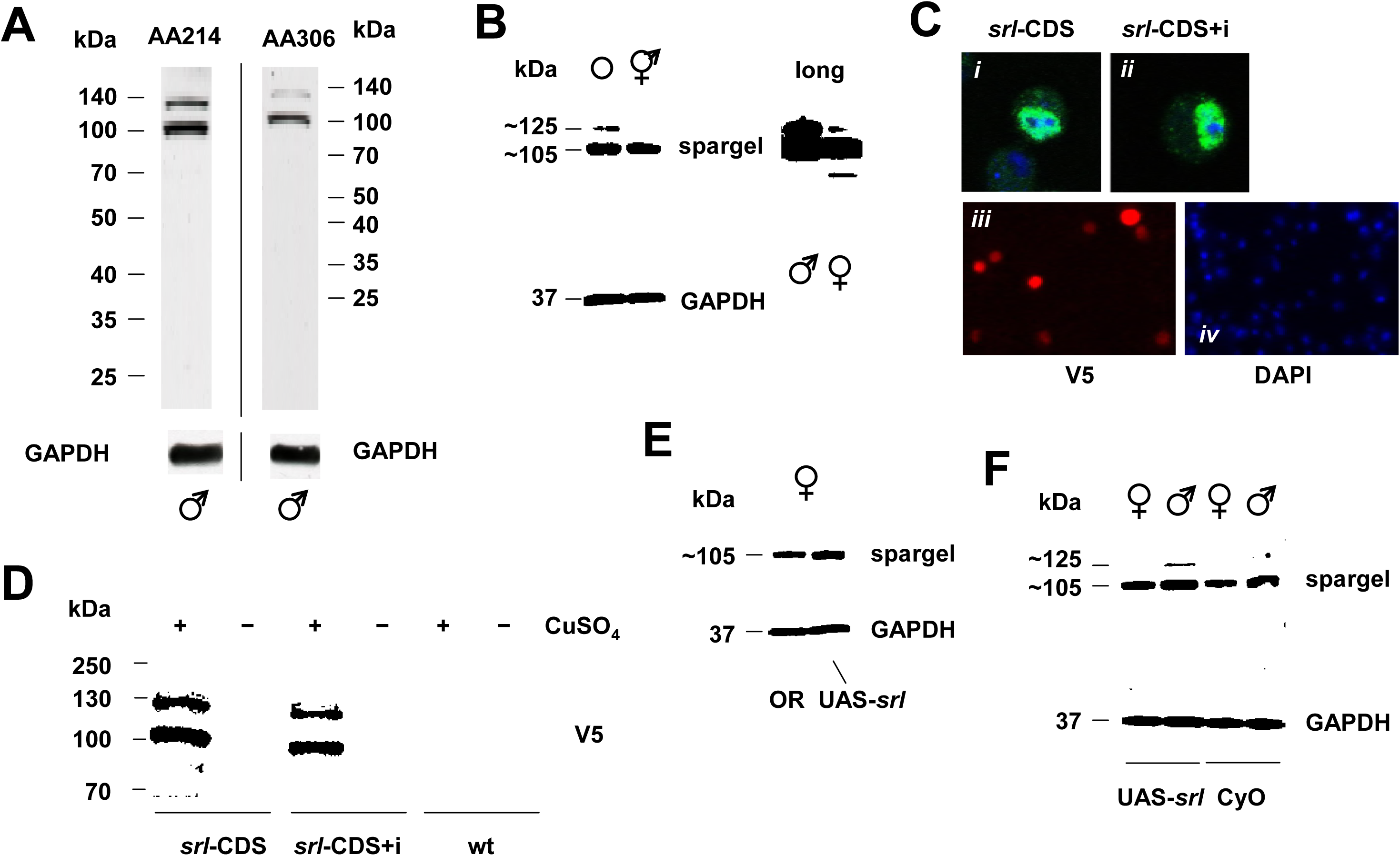
*srl* overexpression at the protein level is modest. (A, B, E, F) Western blots of protein extracts from *Drosophila* adult of the sex and genotype indicated, probed with customized *srl*-directed antibodies AA214 (A, left-hand panel, B, E and F) or AA306 (A, right-hand panel), and with antibody against GAPDH as loading control. Sizes of molecular weight markers in kDa shown in (A), or used to extrapolate sizes of major bands detected in (B, E and F). Note longer exposure of same blot, in right-hand panel of (B). (C) Immunocytochemistry and (D) Western blot, using V5 antibody, on cells transfected with Cu-inducible *srl*-expressing constructs (*srl*-CDS with coding sequence only, *srl*-CDS+i with intron); in (C) panels (i) and (ii) show single cells at high magnification, merged antibody and DAPI, showing ‘speckled’ nuclear localization similar to that observed by Mukherjee & Duttaroy (2013) using *srl*-GFP; panels (iii) and (iv) probed with V5 antibody or DAPI, showing successful transient transfection.

### *srl* overexpression has no systematic effects on *tko*^*25t*^ phenotype

To clarify the effects of *srl* overexpression on the phenotype of *tko*^*25t*^ we conducted a number of tests in which we varied the overexpression construct used, and the genetic background. Using the *srl*^*GR*^ construct we recorded a small decrease in the developmental delay of *tko*^*25t*^ flies modestly overexpressing *srl* (Fig. 3A). However, this was influenced by the presence of the *da*GAL4 driver, since the eclosion day of *tko*^*25t*^ flies lacking both *da*GAL4 and the *srl*^*GR*^ construct was not significantly different from that of flies endowed with both. Furthermore, although the alleviation of developmental delay was significant in this first experiment, as inferred previously (Chen et al., 2012), it was not seen in any of three repeats of the experiment (e.g. the one shown in Fig. 3B). There was also no significant difference in eclosion time between *tko*^*25t*^ flies homozygous for the *srl*^*GR*^ construct and *tko*^*25t*^ controls, in either sex (Fig. 3C). Furthermore, hemizygosity for the extra copy of *srl* produced no rescue of bang-sensitivity (Fig. 3D). More substantial overexpression of *srl* driven by *da*GAL4 using the UAS-*srl* construct did not alleviate developmental delay; rather there was a trend towards a slight deterioration, although this was significant only in one repeat of the experiment, in males only, as shown (Fig. 3E).

**Figure 3:**
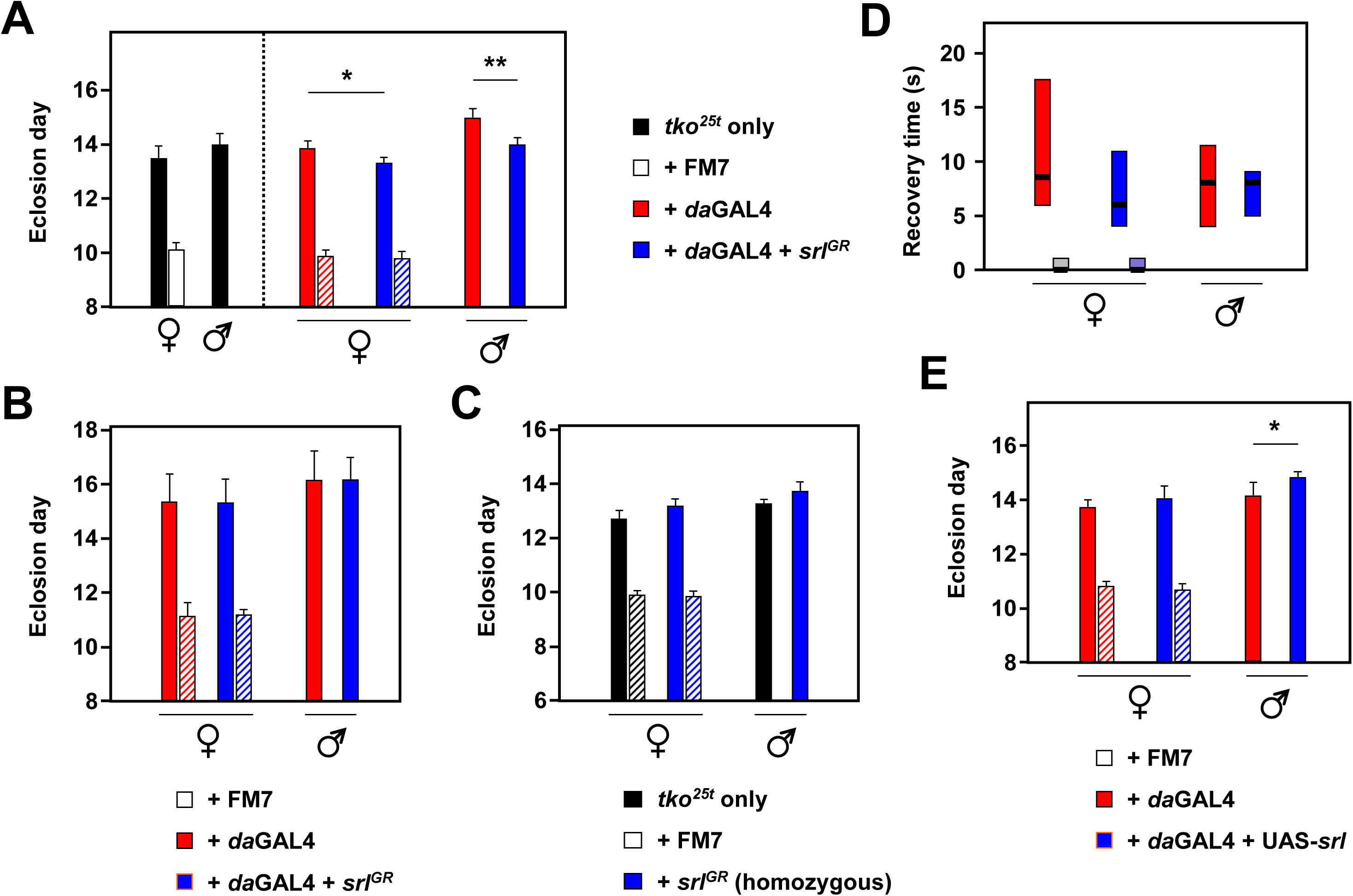
*srl* overexpression does not modify *tko*^*25t*^ phenotype. (A-C, E) Times to eclosion (means ± SD) and (D) bang sensitivity (box plot, 1st to 3rd quartiles, medians as thick black bars) of flies of the indicated genotypes and sex. Dashed vertical line in (A) separates the experimental and ‘*tko*^*25t*^ only’ control (i.e. lacking both *da*GAL4 and *srl*^*GR*^) crosses conducted in parallel (similar controls were implemented routinely but are omitted from the other panels for clarity). (A) and (B) show repeat experiments, in which *tko*^*25t*^; *da*GAL4 males were crossed to *tko*^*25t*^ / FM7 females either with or without *srl*^*GR*^ as shown, which applies also to (D). In (C), crosses for *tko*^*25t*^ alone or in combination with homozygous *srl*^*GR*^ were conducted in parallel, without the presence of *da*GAL4. In (E) progeny carry *da*GAL4 and either UAS-*srl* or CyO from a single cross. Asterisks denote statistically significant differences between progeny flies of a given sex and *tko* genotype, either with or without the presence of an *srl* overexpression construct (Student’s *t* test; *, ** – *p* < 0.05 or 0.01, respectively).

### *srl* expression is not altered by diet during development

Despite these findings, we considered the possibility that the decreased expression of *srl* in *tko*^*25t*^ mimics a response to dietary restriction, resulting in a decreased rate of larval growth. Mitochondrial dysfunction should result in a limitation on energy supply and/or biosynthetic capacity that could activate a growth inhibitory response that would normally be produced by nutritional limitation. *tko*^*25t*^ larvae have decreased levels of ATP, NADPH and the key regulator of cytosolic translation, S6K (Kemppainen et al., 2016). Previously we observed that the developmental delay of *tko*^*25t*^ is influenced by diet: specifically, that media with a low sugar content partially alleviate the phenotype (Kemppainen et al., 2016). We therefore investigated whether culturing *tko*^*25t*^ and wild-type control flies under different nutritional conditions affected *srl* expression in a way that correlated with the phenotypic effect on growth. For this, we compared flies grown on standard high-sugar medium, containing complex dietary additives, with those grown on a minimal medium containing only agar and (10%) yeast. As previously, the low-sugar minimal medium partially accelerated the development of *tko*^*25t*^ flies (Fig. 4A), whilst at the same time retarding that of controls (Fig. 4A, 4B). However, diet-induced effects on the expression of *srl* were minimal. *srl* expression in control (wild-type Oregon R) L3 larvae of both sexes was slightly decreased in minimal medium compared with high-sugar medium (Fig. 4C, 4D), although this was not statistically significant in all experiments (e.g. Fig. 4C, -hand panel). *srl* expression in *tko*^*25t*^ larvae (Fig. 4C, right-hand panel) was lower than in controls by approximately the same factor as in adults, but was unaffected by the different culture media, as was *srl* expression in *tko*^*25t*^ adults (Fig. 4D).

**Figure 4:**
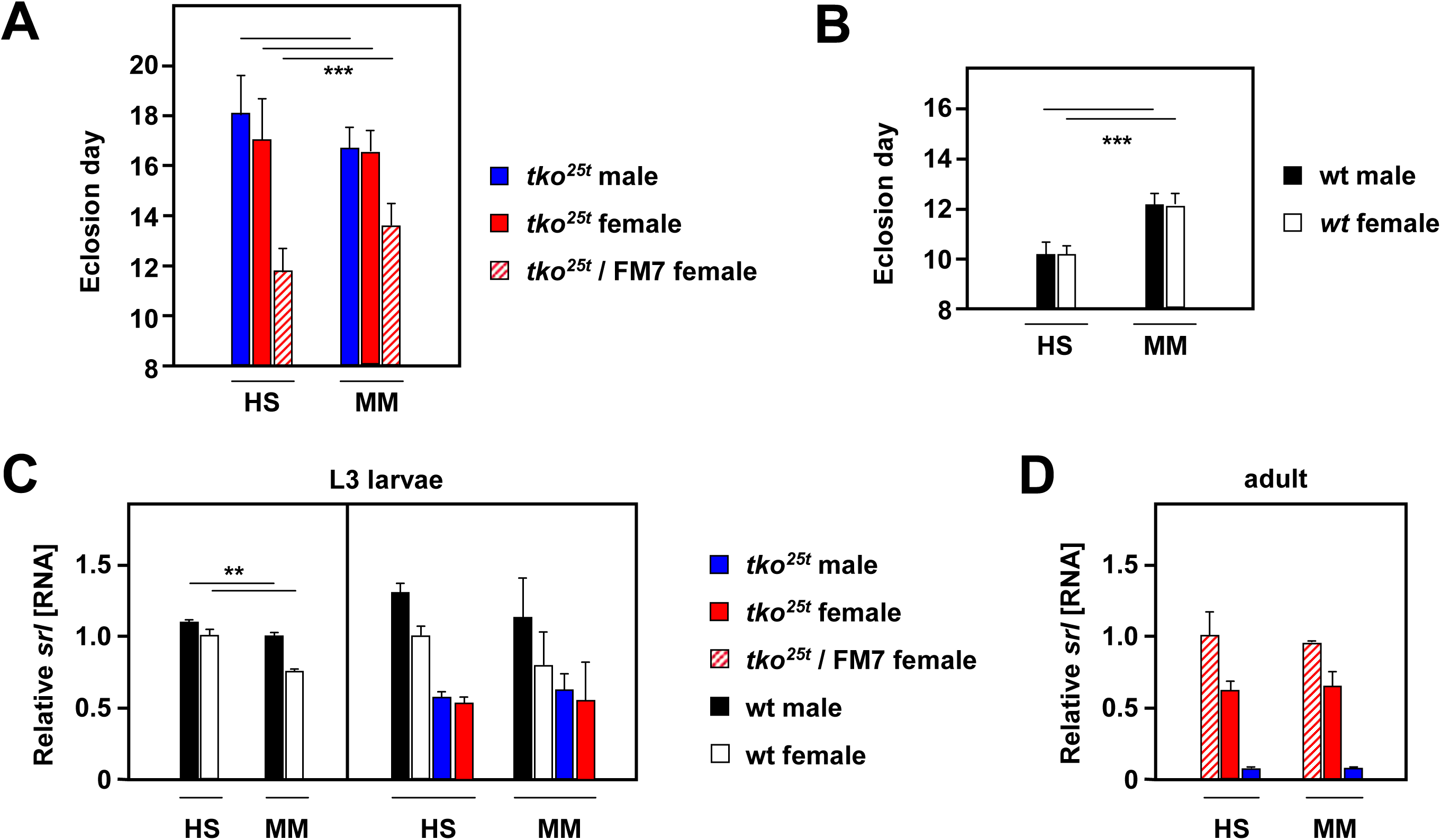
*srl* expression does not correlate with growth rate on different media. (A, B) Means ± SD of times to eclosion of flies of the indicated genotypes and sex, on different media: HS – standard high-sugar medium, MM – minimal medium. wt – wild-type Oregon R. Asterisks (***) denote statistically significant differences between indicated classes of flies of a given sex and genotype, on the different media (Student’s *t* test, *p* < 0.001). Flies grown in (A) bottles, n = 739 individuals, (B) replicate vials, n = 310 flies. (C, D) qRT-PCR measurements of *srl* RNA in L3 larvae and adult flies of the indicated genotypes and sex, on the different media, normalized to values for (C) Oregon R wild-type (wt) females or (D) *tko*^*25t*^ / FM7 heterozygous females. Vertical bar in (C) divides data sets for two separate experiments. Asterisks (**) denote statistically significant differences between indicated classes of flies of a given sex and genotype, on the different media: Student’s *t* test, *p* < 0.01, left-hand part of (C). Note, however, that comparison of values for the equivalent classes in the experiment shown in the right-hand part of (C) gave no significant differences.

### Overexpression of *srl* has no systematic effects on genes related to core mitochondrial functions

Despite the fact that *srl* overexpression had no impact on the *tko*^*25t*^ phenotype, we explored whether such overexpression nevertheless influenced the level of transcripts related to core functions of mitochondria, specifically of mtDNA, nuclear-coded OXPHOS subunits and the major nucleoid protein TFAM (Fig. 5). With the exception of TFAM, all genes studied showed a similar profile of expression in the different strains tested, with higher relative expression in males, higher expression in the *tko*^*25t*^ background, including *tko*^*25t*^ heterozygotes over the FM7 balancer, but which was attenuated by the *da*GAL4 driver and attenuated slightly further by UAS-*srl*. These observations are consistent with expression levels being determined by genetic background, possibly by an effect on the RpL32 reference transcript, rather than by *srl* expression, which followed a different pattern (Fig. 1B). They therefore provide no support for any enhancing effect of *srl*. In the case of TFAM, expression was slightly lower in males than in females, and was little affected by *da*GAL4 or UAS-*srl* (Fig. 5, top right). Note that *srl* overexpression was verified (Fig. 1B) in the same RNA samples.

**Figure 5:**
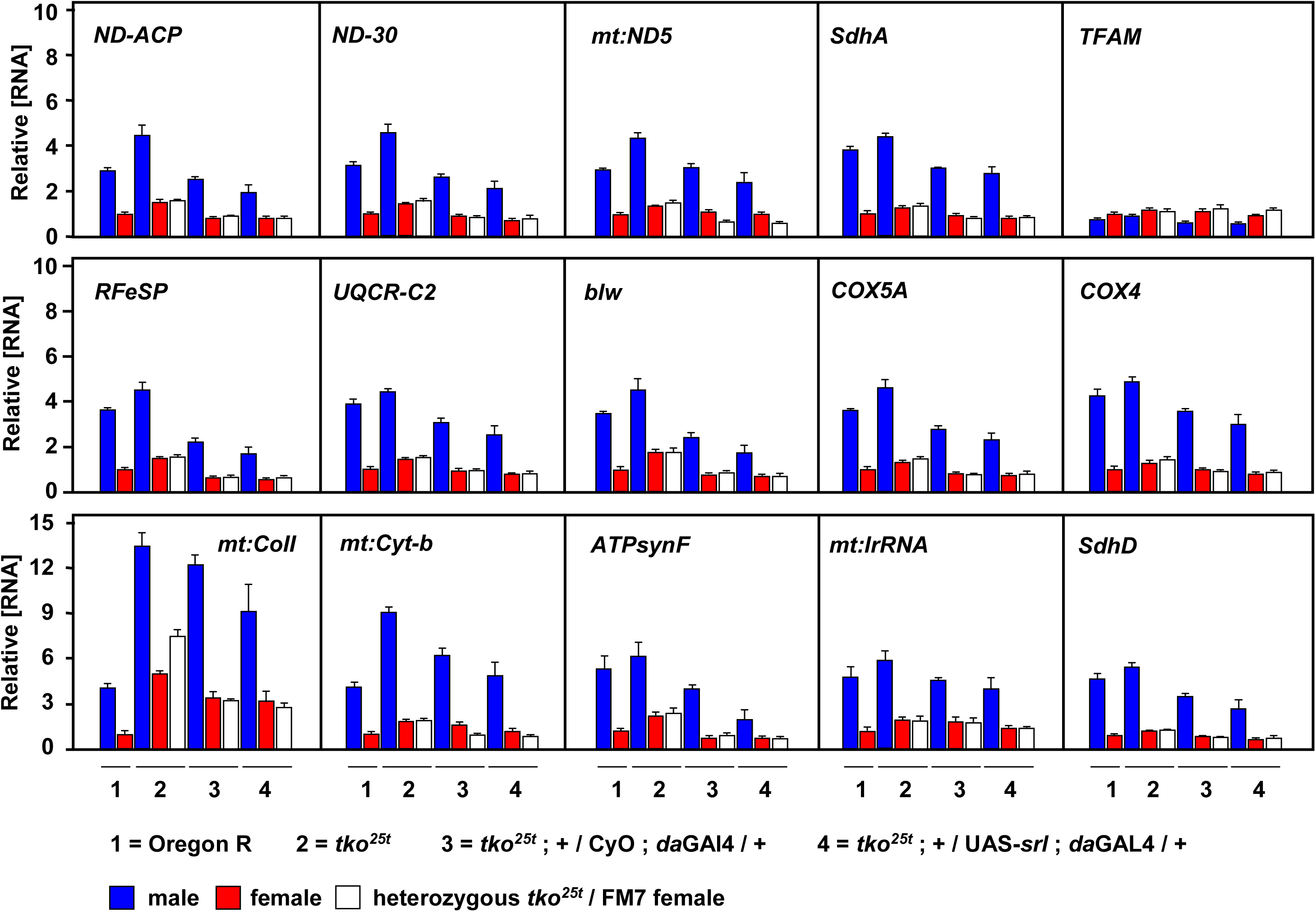
*srl* overexpression does not increase levels of transcripts for core mitochondrial functions. qRT-PCR measurements of RNA levels of the indicated genes (symbols as in flybase.org) in adult flies of the indicated genotypes and sex, normalized to corresponding values for Oregon R (wild-type) females. For clarity, and because expression profiles were qualitatively so similar for all genes studied (except *TFAM*), statistical estimates are omitted.

## DISCUSSION

Previous studies, where the expression of *srl* was downregulated either in the whole fly or in a specific tissue, suggested a global role for *srl* in growth regulation. Whilst those findings are consistent with the proposed role of the PGC1 family of co-activators as ‘master regulators of mitochondrial biogenesis’, other and more general roles in development or gene regulation could not be excluded. Here we addressed the question in an opposite manner, by testing whether overexpression of *srl* was able to compensate the phenotype of *tko*^*25t*^, a mutant with decreased mitochondrial biosynthetic activity. Here too, previous data suggested that *srl* had a modest, beneficial effect.

In the present study, we initially found such a compensatory effect in *tko*^*25t*^ flies endowed with an additional copy of *srl* (Fig. 3A), which boosts expression at the RNA level by a factor of approximately 50% (Fig. 1A). However, the phenotypic effect was not reproduced in several further repeats of the experiment (e.g. Fig. 3B), and a (non-significant) trend in the opposite direction was seen (Fig. 3C), when a further copy of *srl* was added, using a background homozygous for the *srl*^*GR*^-bearing chromosome. *srl*^*GR*^ was also unable to modify the bang-sensitive phenotype of *tko*^*25t*^ (Fig. 3D). Using UAS-*srl* together with *da*GAL4, which gave a much more substantial increase in expression at the RNA level (Fig. 1B), though only a modest effect at the protein level (Fig. 2E, 2F), there was again no alleviation of developmental delay, and even a slight exacerbation (Fig. 3E). Although *srl* mRNA was decreased in *tko*^*25t*^ larvae or adult females compared with controls (Fig. 1C, 1D, 4D), this effect was independent of the growth medium (Fig. 4), and overexpression of *srl* also has no systematic effect on transcripts of genes connected with mitochondrial activity in *tko*^*25t*^. There are several possible, non-exclusive explanations for this essentially negative result that need to be considered.

### Translational regulation

The first possibility is that *srl* is translationally regulated, as suggested by the lack of congruence between RNA and protein analyses. If translational regulation is specific for certain tissues or developmental stages, it may negate overexpression. Although long assumed to be of minor importance, differential regulation at the level of translation is well established (see recent reviews by Zhao et al., 2019; Shi & Barna, 2015), applies to mitochondrial biogenesis itself (Zhang and Xu, 2016), and is prominent in early development (Winata & Korzh, 2018; Barckmann & Simonelig, 2015), especially in *Drosophila*, where it plays roles in axial specification and other processes (Wilhelm & Smibert, 2005; Kugler & Lasko, 2009). Translational regulation is a cardinal feature of the Integrated Stress Response (Ryoo & Vasudevan, 2017), which can be activated by mitochondrial dysfunction.

### Post-translational regulation

The second idea is that *srl* is post-translationally regulated and overexpression is ineffective because another regulatory component of its activity is limiting. Post-translational regulation by a myriad of mechanisms is well documented in diverse contexts, and includes protein phosphorylation (Johnson, 2009), acetylation (Narita et al., 2018), methylation (Lee et al., 2005), proteolytic cleavage (Klein et al., 2018), ligation to peptide modifiers (Gill, 2004) and differential subcellular localization (Bauer et al., 2015). Almost all of these have been documented in the case of PGC-1α (reviewed by Austin & St-Pierre, 2012) and other spargel homologues in mammals. The antibodies that we generated against spargel detect the same bands on Western blots, corresponding in size with those detected by epitope tagging in S2 cells, although their relative amounts vary between transfected cells and flies, and between males and females. The higher molecular weight band (∼125 kDa) plausibly corresponds with the predicted full-length protein of 119 kDa, though might also be modified. The ∼105 kDa band which predominates may be generated by post-translational cleavage, though could also arise by modification or from initiation at an internal AUG (though none seems to be appropriately located: see NCBI entries AAN13314 and AAN13315). It could also be specified by a splice variant, although the one that has been mapped generates an internal 9 amino acid deletion that seems insufficient to account for the observed mobility difference. This issue needs to be clarified in a future study.

Multiple, differentially expressed polypeptides are specified by the mammalian *srl* homologue PGC-1α via alternative promoters and splice-variants, each having distinct proposed functions (Martinez-Redondo et al., 2015), and the same applies to the related genes PGC-1β (Meirhaeghe et al, 2003) and to PPRC1 (predicted from genomic annotation). This complexity makes any extrapolation from the mammalian context hazardous. In theory, the processing that generates the ∼105 kDa spargel polypeptide could activate or inactivate it, or modify its targets, intracellular localization or stability.

### *tko*^*25t*^ signaling

A third possible explanation is that *tko*^*25t*^ elicits a growth inhibitory signal that overrides any effect of *srl*. Such signaling may attune development to the available resources and to the fly’s capacity to use them, enabling the eventual eclosion of a largely normal adult. Although not yet elucidated in detail, these processes effectively coordinate cytosolic with mitochondrial protein synthesis and to the availability of key metabolites. A strong candidate for involvement is ribosomal protein S6 kinase (S6K), whose phosphorylation status responds to signaling via several intersecting pathways such as mTOR (Magnuson et al., 2012), Insulin/Akt (Manning, 2004) and AMPK (Mihaylova & Shaw, 2011), to regulate translation. S6K protein was previously found to be decreased in *tko*^*25t*^ (Kemppainen et al., 2016). Although an enhanced program of mitochondrial biogenesis might be considered one way to correct the metabolic insufficiency of *tko*^*25t*^, the regulation of genes involved in this process is not a global feature of the changes in mRNA expression that occur in *tko*^*25t*^ adults (Fernandez-Ayala et al., 2010), or larvae grown on either high- or zero-sugar medium (Kemppainen et al., 2016). Instead, such changes mainly concern cytosolic translation and protein secretion. Global regulation of translation and growth may therefore mask any effect of *srl* overexpression.

In contradiction to this idea, Mukherjee & Duttaroy (2013) found that *srl* can partially over-ride defects in cell growth mediated by defective insulin/mTOR signaling and that mutants in S6K can be rescued by *srl* overexpression. However, such effects may also be tissue-specific. Alternatively, S6K downregulation may not be instrumental in *tko*^*25t*^, or may be redundant with other growth-limiting pathways.

### A different role for *spargel*

Although *srl* may impact the readout from nutritional signaling, this could be unrelated to mitochondrial biogenesis. A wealth of evidence supports the idea that PGC-1α and its homologues act in mammals to facilitate the transcription of genes that are needed for such processes as adaptive thermogenesis in brown fat (Uldry et al., 2006), neuromuscular junction formation and muscle differentiation in general (Lin et al., 2002; Handschin et al., 2007) and hepatic gluconeogenesis (Yoon et al., 2001) as well as the detoxification of oxygen radicals generated as a by-product of mitochondrial metabolism (St-Pierre et al., 2006). All these processes are linked by the requirement for mitochondrial biogenesis, which is widely held to be the canonical role of the PGC1 coactivators. The reciprocal effects of PGC-1α knockout (Lin et al., 2004; Leone et al., 2005) and its targeted overexpression in the mouse (Lehman et al., 2000; Lin et al., 2002) represent the best evidence of such a role.

On the other hand, this paradigm side-steps the fact that, as a coactivator, it acts, by definition, in co-operation with sequence-specific transcription factors which specify the genes to be regulated. In *Drosophila*, the transcriptional targets of srl are not limited to those involved in mitochondrial biogenesis. In the larval fat body, it was found to promote the transcription of many other growth-related genes responsive to insulin signaling (Tiefenböck et al., 2010).

A critical review of other recent studies relating *srl* expression to physiology provides only weak support for a canonical role in mitochondrial biogenesis. For example, Merzetti & Stavely (2015) found that manipulation of *srl* expression in the eye produces a cell-death phenotype, whilst in dopamine neurons it lead to decreased locomotor activity and effects on lifespan, but these do not identify specific mitochondrial targets. Staats et al. (2018) and Wagner et al. (2015) observed an upregulation of *srl* in flies treated with drugs that promote increased lifespan and locomotor performance in males, but such correlative studies do not address its mechanism of action. *srl* knockdown in muscle did produced morphological abnormalities in mitochondria, whilst its overexpression in dopamine neurons compensated for the effects of *parkin* deficiency (Ng et al., 2017). However, these findings relied on a single (unspecified) RNAi line and on the same overexpression strain used in our study, and did not document effects on mitochondrial gene expression or activity. They are consistent with a global affect of spargel on transcription. Manipulation of *srl* expression similarly affects diet-induced cardiac dysfunction (Diop et al., 2015), but no specificity towards mitochondrial activities has been documented. More convincingly, Rera et al. (2011) reported an increase in mitochondrial markers in flies globally overexpressing *srl* using the same strategy as ourselves (Fig. 1B). However, these markers largely reflect the status of muscle, containing by far the greatest concentration of mitochondria in the fly body, so may indicate a general enhancement of muscle formation and differentiation rather than an effect specific to mitochondria. This may also explain the improved locomotor performance of flies overexpressing *srl* (Tinkerhess et al., 2012).

### Issues in fly genetics

Finally we should consider the possibility that our initial finding using *srl*^*GR*^ (Fig. 3A), was valid, but attributable to some other genetic polymorphism in the *srl*^*GR*^ background that was subsequently lost. Although *srl*^*GR*^ was maintained over a balancer, other relevant polymorphisms in the background may have been eliminated during stock maintenance. Our initial finding also indicates a negative effect of the *da*GAL4 driver in this background.

More generally, this study highlights several important limitations in fly genetics. Balancer chromosomes are an almost universal tool, but allow deleterious mutations to accumulate, protected from negative selection. These potentially compromise the reproducibility of findings in general, and the reliability of quantitating mild phenotypes, such developmental delay in *tko*^*^25^t.*^. Uncontrolled environmental variables, short-term genetic drift and statistical fluctuation should also be considered.

Although burdensome, multiple repeat experiments to confirm quantitatively minor phenotypic variations need to be undertaken, even where findings are judged statistically valid, ideally using independently maintained stocks. Moreover, to minimize background effects, especially where balancer chromosomes restrict back-crossing, such findings should preferably be retested in different backgrounds. Such measures are nevertheless much easier to implement and interpret in *Drosophila*, compared with mammalian models where inconsistent or strain-dependent findings abound.

## MATERIALS AND METHODS

### *Drosophila* strains and culture

The *srl*^*GR*^ and UAS-*srl* strains (Tiefenböck et al., 2010), both supplied over a CyO balancer, were a kind gift from Christian Frei (ETH Zürich). The *tko*^*25t*^ strain, originally sourced through Kevin O’Dell (University of Glasgow), was backcrossed into Oregon R background (Toivonen et al., 2001) and maintained long-term in our laboratory over the FM7 balancer. The Oregon R wild-type and *da*GAL4 driver strains were originally obtained from Bloomington Stock Centre, and the *tub*GS driver was the kind gift of Scott Pletcher (University of Michigan). All stocks were maintained at room temperature and grown experimentally in plugged plastic vials at 25 °C on a 12 h light/dark cycle in standard high-sugar medium (HS, Kemppainen et al., 2016) or, where specified in figures, in a minimal medium (MM) consisting of agar, 10% yeast and standard antimicrobial agents (0.1% nipagin and 0.5% propionic acid, Sigma-Aldrich),

### Molecular cloning

Genomic DNA was extracted from adult *Drosophila* and used as a PCR template with chimeric gene-specific primers to amplify *srl* from the start codon up until, but not including, the stop codon. The chimeric primers contained EcoRI and NotI restriction sites for restriction digestion and insertion into the copper-inducible plasmid pMT-V5/HisB (Thermo Fisher Scientific), resulting in the introduction of an in-frame C-terminal V5 epitope tag. A primer deletion strategy was used on this plasmid to create an intronless version of *srl* tagged with V5. Both resulting plasmids were sequence-verified before use in transfections.

### Developmental time and bang-sensitivity assays

Three replicate crosses were set up and tipped five times to fresh vials for egg-laying, as previously (Kemppainen et al., 2009). The mean developmental time to eclosion (at 25°C), as well as bang-sensitivity were measured as described previously (Kemppainen et al., 2009). Unweighted means and standard deviations of eclosion day for each sex and inferred genotype were then computed for each cross, and used in statistical analyses, generally applying Student’s *t* test (unpaired, two-tailed) to compare the mean eclosion day of flies of a given sex and genotype, with and without the expression of a given *srl* overexpression construct. For bang-sensitivity, medians and quartiles of recovery time for flies of a given sex and genotype were plotted in a box-plot format.

### RNA analysis

Total RNA was extracted from batches of ten 2 day-old flies and from L3 (wandering stage) larvae using a homogenizing pestle and trizol reagent as previously described (Kemppainen et al., 2016). cDNA was synthesized using the High-capacity cDNA Reverse Transcription Kit (ThermoFisher Scientific) according to manufacturer’s instructions. Expression levels were determined by qRT-PCR using Applied Biosystems StepOnePlus™ Real-Time PCR System with Fast SYBR™ Green Master Mix kit (Applied Biosystems) using, as template, 2 μl of cDNA product diluted 10-fold, in a 20 μl reaction, together with 500 nM of each gene-specific primer pair as follows (all given 5’ to 3’, gene symbols in the following list following current practice in flybase.org): *RpL32* (CG7939), TGTGCACCAGGAACTTCTTGAA and AGGCCCAAGATCGTGAAGAA; *ND-ACP* (CG9160), ACAAGATCGATCCCAGCAAG and ATGTCGGCAGGTTTAAGCAG: *ND-30* (CG12079), AAGGCGGATAAGCCCACT and GCAATAAGCACCTCCAGCTC; *mt:ND5* (CG34083), GGGTGAGATGGTTTAGGACTTG and AAGCTACATCCCCAATTCGAT; *SdhA* (CG17246), CATGTACGACACGGTCAAGG and CCTTGCCGAACTTCAGACTC; *TFAM* (CG4217), AACCGCTGACTCCCTACTTTC and CGACGGTGGTAATCTGGGG; *RFeSp* (CG7361), GGGCAAGTCGGTTACTTTCA and GCAGTAGTAGCCACCCCAGT; *UQCR-C2* (CG4169), GAGGAACGCGCCATTGAG and ACGTAGTGCAGCAGGCTCTC; *Blw* (CG3612), GACTGGTAAGACCGCTCTGG and GGCCAAGTACTGCAGAGGAG; *COX5A* (CG14724), AGGAGTTCGACAAGCGCTAC and ATAGAGGGTGGCCTTTTGGT; *COX4* (CG10664), TCTTCGTGTACGATGAGCTG and GGTTGATTTCCAGGTCGATG; *mt:CoII* (CG34069), AAAGTTGACGGTACACCTGGA and TGATTAGCTCCACAGATTTC; *mt:Cyt-b* (CG34090), GAAAATTCCGAGGGATTCAA and AACTGGTCGAGCTCCAATTC; *ATPsynF* (CG4692), CTACGGCAAAGCCGATGT and CGCTTTGGGAACACGTACT; *mt:lrRNA* (CR34094), ACCTGGCTTACACCGGTTTG and GGGTGTAGCCGTTCAAATTT; *SdhD* (CG10219), GTTGCAATGCCGCAAATCT and GCCACCAGGGTGGAGTAG; *srl* (CG9809), GGAGGAAGACGTGCCTTCTG and TACATTCGGTGCTGGTGCTT. Mean values were normalized first against that for *RpL32* and then against an arbitrary standard, wild-type (Oregon R) adult females, except where stated.

### Protein analysis

Batches of ten 2 day-old adult flies were crushed in 100 µl of lysis buffer (0.3% SDS in PBS plus one EDTA-free cOmplete™ Protease Inhibitor Cocktail tablet, Roche), incubated for 15 min and centrifuged at 15,000 *g*_*max*_ for 10 min (all manipulations at room temperature). Supernatants were decanted and protein concentrations determined by the Bradford assay. Aliquots of 50 μg protein in SDS-PAGE sample buffer containing 0.2 M dithiothreitol were heat-denatured for 5 min at 95 °C then electrophoresed on AnyKD midi criterion™ gels (Bio-Rad) in ProSieve™ EX running buffer (Lonza). Transfer to Nitrocellulose membrane (Perkin-Elmer) was performed using ProSieve™ EX transfer buffer (Lonza). Membranes were blocked in 5% nonfat milk in PBS-0.05% Tween (Medicago) for 30 min at room temperature, with gentle agitation. Primary antibody diluted in the same buffer was added and reacted at 4 °C overnight. After three 10 min washes, secondary antibody was added in the same buffer containing 5% nonfat milk in for a further 2 h. Membranes were washed twice for 10 min in PBS-0.05% Tween and then for a final 10 min in PBS. Primary antibodies and dilutions were as follows: Srl214AA (against peptide CFDLADFITKDDFAENL) and Srl306AA (against peptide CPAKMGQTPDELRYVDNVKA), custom rabbit polyclonal antibodies (21st Century Biochemicals, both 1:5000), GAPDH (Everest Biotech EB06377, goat polyclonal, 1:5000), anti-V5 (ThermoFisher Scientific, mouse monoclonal #R96025, 1:10000). Appropriate HRP-conjugated secondary antibodies (Vector Laboratories) were used at 1:5,000). 5 ml of Luminata™ Crescendo Western HRP substrate solution (Merck) was added for 5min before imaging with a Bio-Rad imager.

### Transfections and immunocytochemistry

Transfection and induction of S2 cells with V5-tagged *srl* constructs and subsequent staining for imaging was performed as previously (González de Cózar et al., 2019). The primary antibody used was mouse anti-V5 (Life Technologies) along with the corresponding Alexa Fluor® 488 or Alexa Fluor® 647 secondary antibodies (Abcam), with image acquisition by confocal microscopy.

### Image processing

Images have been cropped and/or rotated for clarity and optimized for contrast and brightness, but without other manipulations.

## ACKNOWLEDGEMENTS

We thank Tea Tuomela and Eveliina Teeri for technical assistance.

## AUTHOR CONTRIBUTIONS

HTJ and JG jointly conceived the study. JG performed the experiments and analyses. HTJ supervised the work, compiled the figures and drafted the manuscript, which was finalized by both authors.

